# The effect of response biases on resolution thresholds of Sloan letters in central and paracentral vision

**DOI:** 10.1101/2020.09.04.283119

**Authors:** Hatem Barhoom, Mahesh R Joshi, Gunnar Schmidtmann

## Abstract

**Purpose:** Letter resolution depends on three factors: perceivability, response bias, and similarity. The aim of the present study was to investigate the effect of response bias (the sensory-independent factor) on resolution thresholds of Sloan letters in central and paracentral vision.

**Methods:** Nine subjects with normal ocular health were recruited for this study. Using the method of constant stimuli, individual Sloan letters resolution thresholds were measured at 0° (central) and at ±3° eccentricity along the vertical meridian of the visual field. Response biases and letter similarities were computed using Luce’s choice model.

**Results:** Results showed that the differences in resolution thresholds of individual Sloan letters were significant at the central (F (9, 80) =5.02, *p*<.001), the upper (χ2 (9) = 50.38, *p*<.05) and the lower (χ2 (9) = 56.32, *p*<.05) visual field locations. Unlike letter similarity measures, response biases showed significant correlations to the differences in thresholds at the central (r = −0.83, *p*<.05), the upper (r = - 0.73, *p*<.05) and the lower (r = −0.70, *p*<.05) visual field locations.

**Conclusions:** Our findings suggest that response biases have a significant effect on resolution of Sloan letters that could result in overestimating resolution acuity in central and paracentral visual field locations.

## Introduction

Visual acuity is the ability of the visual system to discern the smallest details of an object, typically measured as the minimum angle of resolution (i.e. detection / resolution threshold) and is of high clinical importance. Clinically, many stimuli or optotypes, such as the Tumbling E, Landolt C and alphanumeric characters have been employed to measure visual acuity (Kniestedt & Stamper, 2003). Although the Landolt C is internationally regarded as the reference optotype (Sloan, 1959; Treacy, Hurst et al., 2015), letters are used in many visual acuity charts, because they are intuitive and easy to use in clinical settings. Furthermore, employing a variety of letters (e.g. Sloan letters) reduces the guessing rate associated with forced choice tests (Pelli & Robson, 1991).

In general, letter identification accuracy is influenced by three factors: (i) perceivability, (ii) response bias and (iii) similarity (Mueller & Weidemann, 2012). Perceivability is a measure of how legible the letter is depending solely on the characteristics of the letters, such as the letter size, contrast or shape. The response bias is defined as the tendency of favouring one response over the other alternatives (Macmillan & Creelman, 1990), and similarity is defined as the confusion in letter perception which arises among certain letters. In other words, letter recognition, i.e. the letter detection / resolution threshold, could be affected by changing the amount or the type of the sensory input, e.g. size and contrast (perceivability), the bias towards certain letters in case of uncertainty (response biases), and the confusion between ‘similar’ letters such as for instance **C** and **O** (similarity). Note that from these definitions it is well understood that response biases, unlike perceivability and letter similarities, are independent from the sensory inputs of the stimulus.

Unlike other common letter stimuli, Sloan letters have been adopted in the design of many letter charts (e.g. Early Treatment Diabetic Retinopathy Study; ETDRS chart), because their average legibility, determined by the letter identification accuracy is similar to the difficulty in resolution of the Landolt C (Sloan, 1959; Treacy, Hurst et al., 2015). It has been shown that Sloan letters have different relative legibility of the individual letters at the fovea where the letter similarities are the major source of errors in threshold determination (McMonnies & Ho, 1996; Reich & Bedell 2000; Hamm, Yeoman et al., 2018). However, little is known about the effect of response biases on the resolution thresholds of individual Sloan letters. In this study, the aim was to investigate the effect of the response biases on resolution thresholds of individual Sloan letters in central and paracentral locations since the pattern of differences in letter thresholds has been found to be different at the central and paracentral locations (Ludvigh, 1941; Strasburger, Rentschler et al., 2011; Hairol, Abd-Latif et al., 2015).

## Methods

### Participants

Nine naïve subjects (six females and three males, mean age 22.89 ±3.51 (SD) and age range from 19 to 28 years old) with normal ocular health participated in this study. The mean best corrected visual acuity and the mean refractive error (spherical equivalent) were −0.041 ±0.066 logMAR and −2.36 ±2.43 *DS* respectively. All tests were done monocularly (left or right eye, chosen at random), where the fellow eye was occluded using an opaque eye patch. Written informed consent was obtained from all observers, and the study was approved by the University of Plymouth Ethics committee. All experiments were conducted in accordance with the Declaration of Helsinki.

### Apparatus

Stimuli were generated using MATLAB (MathWorks, Natick, Massachusetts, USA) (MATLAB R2016b, MathWorks). The stimuli were presented on a gamma-corrected DELL, P2317H LCD monitor (1920×1080) with a frame rate of 60Hz. Monitor linearization was achieved by adjusting its colour look-up table, resulting in 150 approximately equally 2 spaced grey levels. Observers viewed the targets in a viewing distance of 350 cm, while sitting on chair without using a chin or forehead rest. The examiner guaranteed a constant viewing distance by regular check. At this viewing distance one pixel subtended 0.258 minutes of arc (’) of visual angle. Experiments were carried out under room illumination of 160 lux. The observer responded by calling out the responses which were entered by the experimenter via a standard computer keyboard. This method minimised errors caused by mistyping and improved fixation compliance. Routines from the PsychToolbox were used to present the stimuli (Brainard, 1997; Pelli, 1997; Kleiner, Brainard et al., 2007).

### Stimuli

High contrast Sloan letters were used as stimuli for the experiment (black letters of 2.2 cd/m^2^ on a white background of 215 cd/m^2^, resulting in 99% letter Weber contrast). The variables were letter size, expressed in minutes of arc, subtended by the stroke width of the letter at the viewing distance, and the location of presentation. The letters were presented centrally and at paracentral locations along the vertical meridian at an eccentricity of 3° in the upper and lower visual field (Figure 1a). Ten standard Sloan letters: **C**, **D**, **H**, **K**, **N**, **O**, **R**, **S**, **V**, **Z** were used. Each Sloan letter is designed so that its height is equal to its width and five times the stroke width (Figure 1b). Six different letter sizes (spaced logarithmically) were tested; 0.3’, 0.44’, 0.64’, 0.94’, 1.37’, and 2’ for central presentations and 0.5’, 0.79’, 1.26’, 1.99’, 3.15’ and 5’ for paracentral presentations.

**Figure 1.**
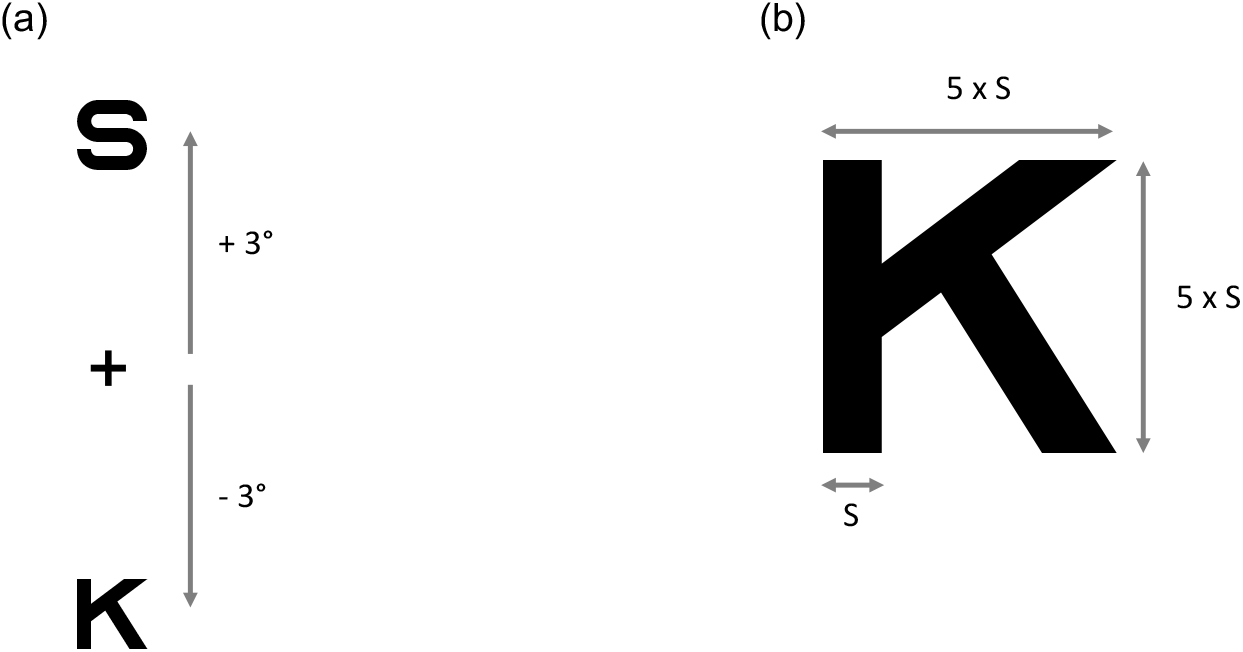
(a) Sloan letters presented centrally, or along the vertical meridian at an eccentricity of 3° in the upper and lower visual field. The Sloan letters **S** and **K** are shown for illustration purposes (not to scale). (b) shows the dimensions of the Sloan letters, exemplified by the letter **K**. The stroke width S = 1/5 of the letter’s height. The height and the width of the letter are equal.

### Procedure

The method of constant stimuli was used for all experiments in this study. The Sloan letters were presented randomly across the three locations, so that at each location each letter was presented 10 times for six different letter sizes. The presentation time was 250 ms and presentations were accompanied by an auditory signal. The task was to recognise the presented letter and to report it verbally. During the experiment, the subject was asked to fixate on a fixation cross (Dimensions: length/width 1.55 min arc, stroke width 0.036 min arc) presented in the middle of the screen. Subjects were encouraged to guess when uncertain about the letter. Only choices of the 10 Sloan letters were accepted. When the observer responded with a non-Sloan letter, the experimenter prompted for a second response from the Sloan set. Each subject completed 1800 trials for the full experiment (six letter sizes × three locations ×10 Sloan letters ×10 presentations per letter).

### Analysis

Data analysis was performed using MATLAB. Routines from the Palamedes Toolbox (Prins & Kingdom, 2018) were employed to fit individual psychometric functions. The data (percent correct vs. letter size) were fit with Gumbel (Log-Weibull) functions (expressed by the general formula (Eq.1)) for each of the 10 letters at three locations.

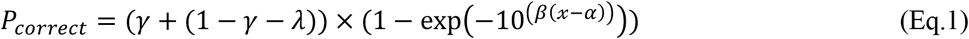

where *γ* is the guessing rate, *λ* is the lapse rate, *x* is the letter size (log visual angle), *α* is the threshold and *β* is the slope of the function. With guessing rate of 0.1 (10 letters) and lapse rate of 0.02 (naïve subjects), the threshold *α* was defined as *x* yielding 65.6% correct responses, according to the following equation.

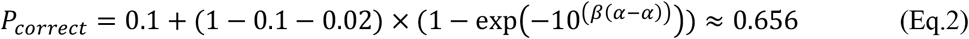

## Results

The mean thresholds across 10 Sloan letters (±SD) were 0.059’ ±0.074, 0.425’ ±0.145 and 0.445’ ±0.141 log visual angle at the central location, upper visual field (3°) and lower visual field (3°), respectively. Paired sample t-test results showed no significant difference between the mean thresholds at upper (0.425’ ±0.145) and lower visual field location (0.445’ ±0.141) (t (8) = −0.616, *p*=.55). However, the mean thresholds at the upper visual field (t (8) = 6.95, *p*<.001) and lower visual field (t (8) = −7.71, *p*<.001) were significantly higher than the central location. The mean thresholds for individual letters at each location are shown in Table 1. One-way ANOVA tests showed statistically significant differences between the mean thresholds of individual letters at the central location (F (9, 80) = 5.02, *p*<.001) with highest and lowest thresholds for **V** = 0.14 ±0.08 and **K** = −0.01 ±0.11 log visual angle (minutes of arc) respectively. Additionally, Kruskal-Wallis H tests revealed statistically significant differences in the estimated thresholds of individual letters at the upper (χ^2^ (9) = 50.38, *p*<. 05) (with highest and lowest thresholds for **V** = 0.61±0.12 and **N** = 0.22 ±0.07 log visual angle (minutes of arc) respectively) and lower (χ^2^ (9) = 56.32, *p*<.05) (χ^2^ (9) = 50.38, *p*<.05) (with highest and lowest thresholds for **V** = 0.66 ±0.28 and **S** = 0.20 ±0.07 log visual angle (minutes of arc) respectively) visual field locations.

**Table 1.** shows the mean individual resolution thresholds for letters in log visual angle (minutes of arc) at central and paracentral locations.

| | Central | Upper field ($3^\circ$) | Lower field ($3^\circ$) |
| --- | --- | --- | --- |
| **C** | $0.05' \pm 0.07$ | $0.40' \pm 0.09$ | $0.45' \pm 0.09$ |
| **D** | $0.04' \pm 0.10$ | $0.35' \pm 0.08$ | $0.58' \pm 0.12$ |
| **H** | $0.07' \pm 0.09$ | $0.24' \pm 0.10$ | $0.33' \pm 0.09$ |
| **K** | $-0.01' \pm 0.11$ | $0.44' \pm 0.10$ | $0.45' \pm 0.08$ |
| **N** | $0.11' \pm 0.05$ | $0.22' \pm 0.07$ | $0.29' \pm 0.05$ |
| **O** | $-0.05' \pm 0.05$ | $0.48' \pm 0.16$ | $0.46' \pm 0.12$ |
| **R** | $0.01' \pm 0.07$ | $0.41' \pm 0.10$ | $0.33' \pm 0.12$ |
| **S** | $0.02' \pm 0.09$ | $0.25' \pm 0.10$ | $0.20' \pm 0.07$ |
| **V** | $0.14' \pm 0.08$ | $0.61' \pm 0.12$ | $0.66' \pm 0.28$ |
| **Z** | $-0.05' \pm 0.11$ | $0.45' \pm 0.17$ | $0.42' \pm 0.14$ |

For the following analyses, data were pooled across subjects, and confusion matrices for each location were created. Table 2 shows the overall and individual letters thresholds for each location. Figure 2 shows the confusion matrices (presented vs. letter response) for each location. The greyscale illustrates the frequency of letter response, where darker cells show higher frequencies. The diagonal cells represent correct responses, whereas the non-diagonal cells represent incorrect responses.

**Figure 2.**
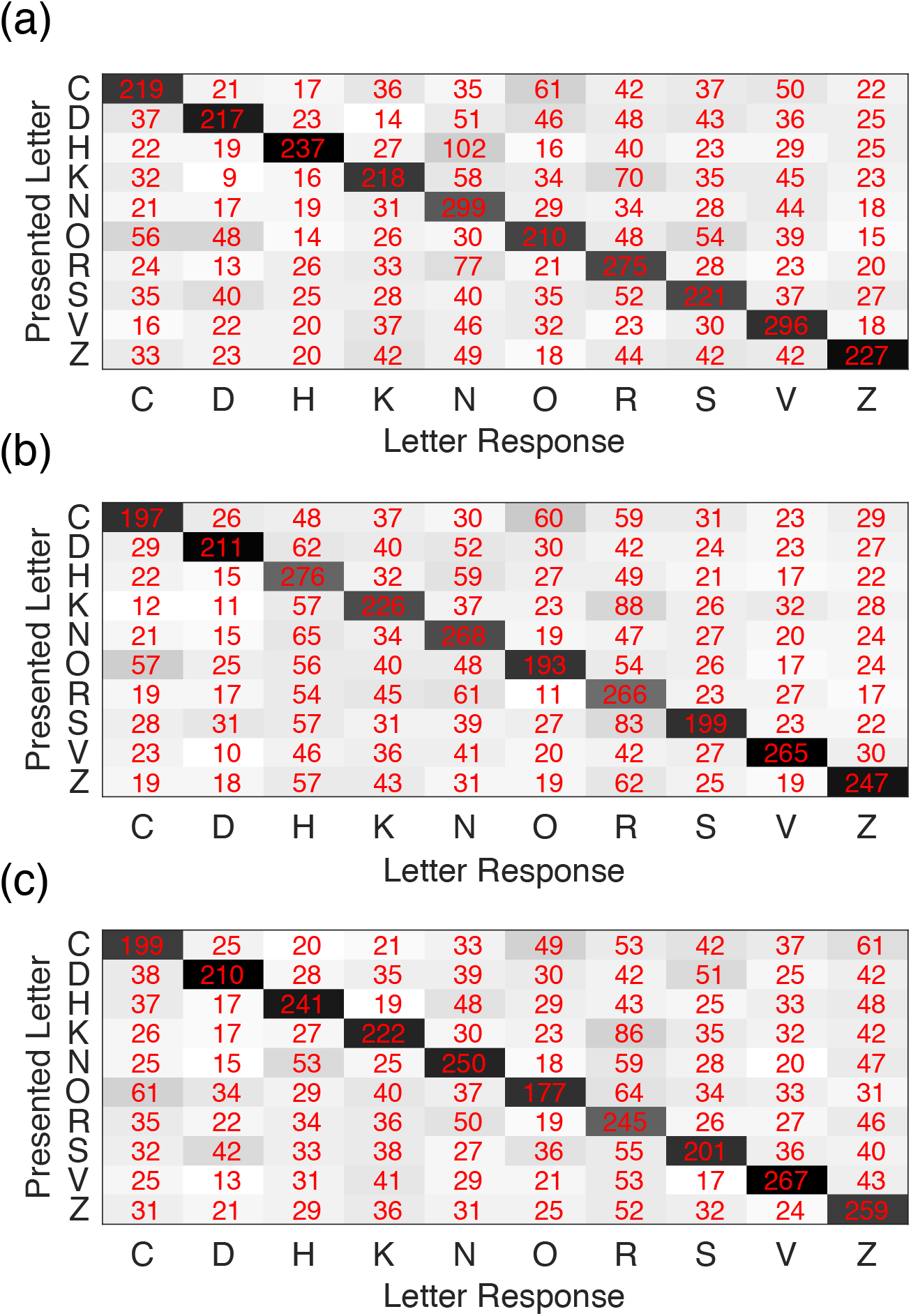
shows the confusion matrices of the three locations (a) central vision, (b) upper visual field (3°) and (c) lower visual field (3°). The number in each cell represents the frequency of the answered letter to the presented letter corresponding to that cell, pooled over subjects and letter sizes. The diagonal cells represent the correct responses, whereas the non-diagonal cells represent incorrect responses.

**Table 2.** shows the overall and the individual resolution thresholds (pooled across subjects) in log visual angle (minutes of arc) for letters at central and paracentral locations.

|  | Central | Upper field (3°) | Lower field (3°) |
| --- | --- | --- | --- |
| Overall | 0.06' | 0.45' | 0.46' |
| **C** | 0.14' | 0.53' | 0.53' |
| **D** | 0.09' | 0.48' | 0.44' |
| **H** | 0.05' | 0.35' | 0.39' |
| **K** | 0.10' | 0.46' | 0.49' |
| **N** | -0.04' | 0.39' | 0.42' |
| **O** | 0.12' | 0.57' | 0.59' |
| **R** | -0.01' | 0.39' | 0.42' |
| **S** | 0.12' | 0.56' | 0.58' |
| **V** | -0.05' | 0.34' | 0.35' |
| **Z** | 0.06' | 0.39' | 0.36' |

### Model

The difference in resolution thresholds between letters can be caused by the difference in the relative legibility of the letters, response biases and/or letter similarities. To investigate the potential effect of these three factors on the letter detection thresholds, response biases and letter similarities were computed using Luce’s choice model (Luce, 1963). This model attempts to disentangle the response factor that is sensory-independent (i.e. response biases towards some letters) from the sensory-dependent response factor (i.e. similarities between certain letters). Here Luce’s model was used to estimate the response biases (expressed in response bias vector) (Eq. 3) and letter similarities (Eq. 4). The model predications are presented as similarity matrix capturing the similarity between each pair of letters parameters. These parameters were calculated from the matrices of the maximum likelihood estimates which resulted from the model fit (see appendix). According to the Luce’s choice model:

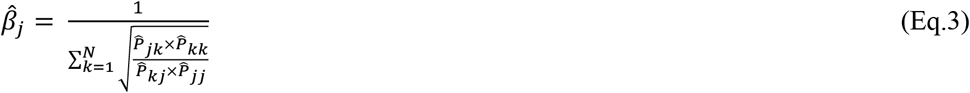

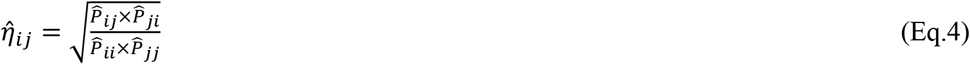

(Eq.4) where the variable *β* in Eq. 3 denotes the response bias parameter for the letter *j. N* is the number of letters (10 letters). *η* is the similarity parameter of each cell between the letter *i* and the letter *j*. 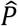 is the expected value in each cell obtained from the maximum likelihood estimates matrix.

Using this method, the most prominent response biases (values higher than the average of *β* parameters) were found to be towards the letters **N**, **R**, **V** at the central location, **H**, **N**, **R** at the upper and **N**, **R**, **V**, **Z** at the lower visual field locations. Figure 3 shows the response biases (*β* parameters in Eq. 3) for each letter depicted for central location (**C** = 0.087, **D** = 0.077, **H** = 0.08, **K** = 0.09, **N** = 0.152, **O** = 0.086, **R** = 0.128, **S** = 0.094, **V** = 0.128, **Z** = 0.077), upper (**C** = 0.073, **D** = 0.069, **H** = 0.146, **K** = 0.104, **N** = 0.13, **O** = 0.074, **R** = 0.144, **S** = 0.077, **V** = 0.092, **Z** = 0.091) and lower (**C** = 0.089, **D** = 0.076, **H** = 0.102, **K** = 0.096, **N** = 0.112, **O** = 0.073, **R** = 0.133, **S** = 0.086, **V** = 0.108, **Z** = 0.126) visual field locations. The letter similarities for letter pairs (*η* parameters in Eq. 4) are shown in Figure 4 as triangular matrices for the central and paracentral locations.

**Figure 3.**
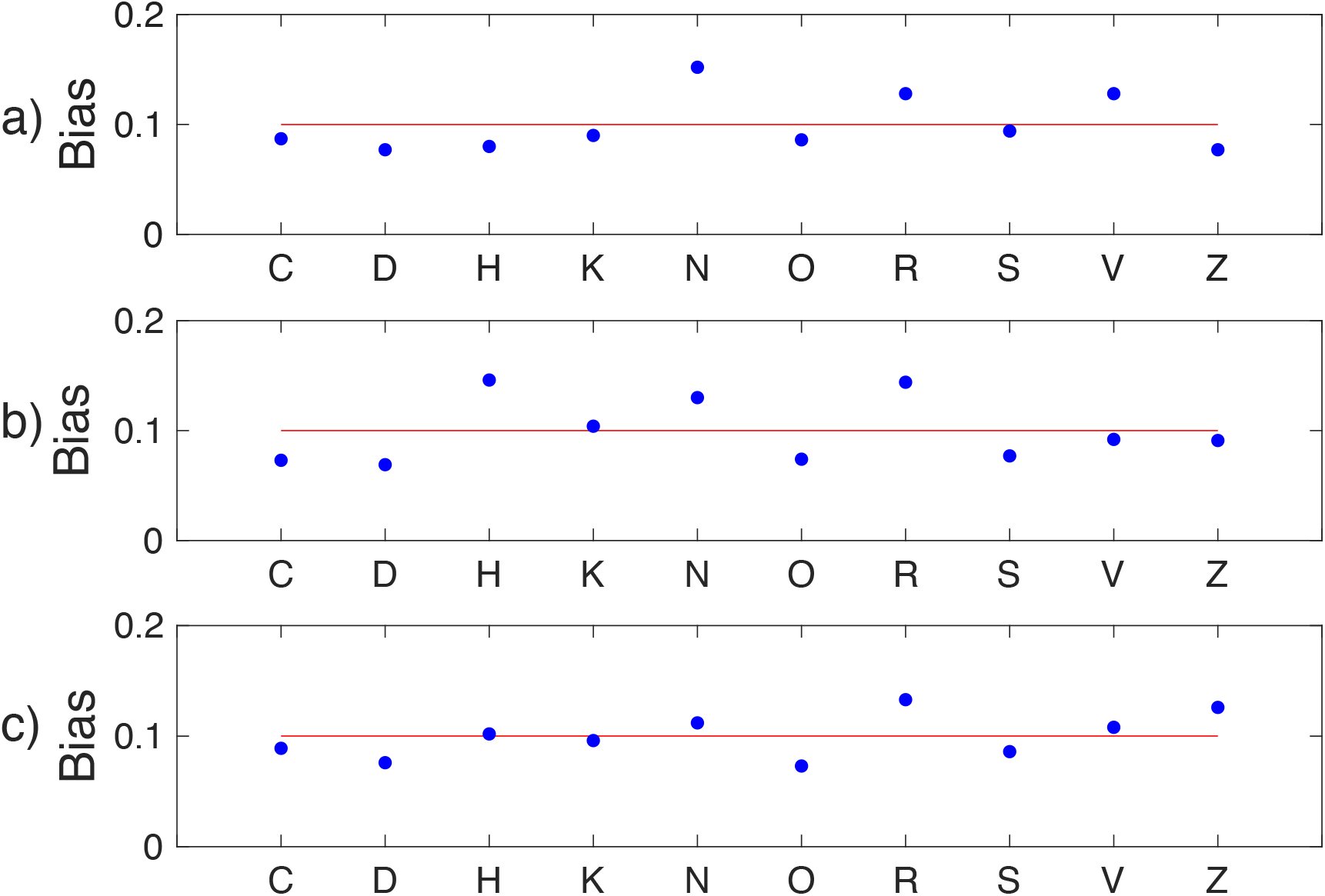
shows the response biases calculated by Luce choice model at the (a) central location, (b) upper (3°) and (c) lower (3°) visual field. The blue dots represent the biases as *β* parameters. The red horizontal line in each plot is the average of *β* parameters.

**Figure 4.**
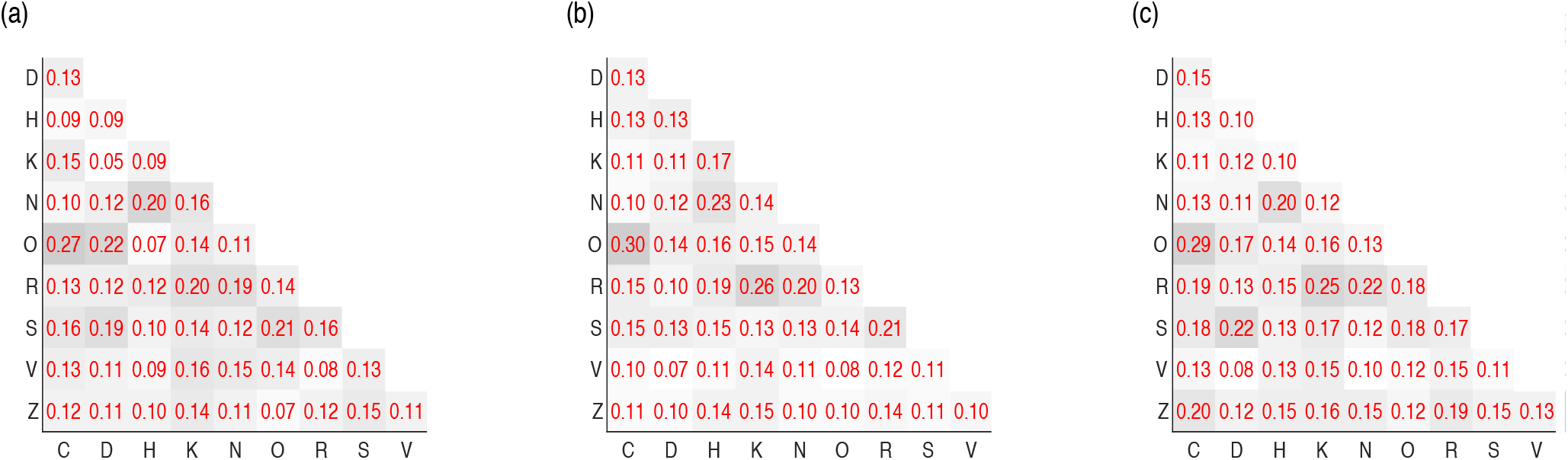
shows the similarity calculated by the Luce choice model at the (a) central, (b) upper (3°) and (c) lower (3°) visual field locations. The numbers in each cell are the similarity parameters (*η*). Darker cells show that the two corresponding letters are perceived to be more similar.

For further analysis, the ‘confusability’ of each letter was calculated as the average of the *η* parameters for the letter and is therefore considered as a measure of the confusability of the letter with the remaining letters. These parameters, *β* and *η* were used to determine the effect on the thresholds differences of letters as discussed in the next paragraph.

To assess the effect of the letter similarities and the response biases on the thresholds we investigated the correlation of the *β* and *η* parameters (expressed as confusability of letters, described above) to the letters’ individual thresholds. Pearson product-moment correlation test showed that the correlation between the confusability and the individual thresholds of letters was not statistically significant at any of the three locations (central: *r* = 0.26, *n* = 10, *p* = .47; upper: *r* = 0.09, *n* = 10, *p* = .8; lower: *r* = 0.52, *n* = 10, *p* = .12). Interestingly, these tests revealed a statistically significant negative correlation between letter resolution and response biases at all three locations (central: *r* = −0.83, *n* = 10,*p* < .05; upper: *r* = −0.73, *n* = 10,*p* < .05; lower: *r* = −0.70, *n* = 10,*p* < .05).

These findings suggest that response biases had a significant effect on the resolution thresholds of the individual letters, while confusability (average letter similarities) did not. In short, lower thresholds were associated with higher response biases.

## Discussion

The aim of this study was to investigate the effect of response biases on the resolution thresholds of individual Sloan letters. Results show that thresholds are statistically significant different between the central and the paracentral visual field locations with no significant difference between the upper (3°) and lower (3°) location. These results are similar to those reported previously. Figure 5 shows the slope of the regression line of letter resolution log thresholds as a function of eccentricity for two previous studies (Ludvigh, 1941; Hairol, 2015) and the current one. A direct comparison of the regression lines of two previous studies and the current one shows a similar slope, but higher thresholds (Figure 5). The higher thresholds in this study could be the result of differences in study design. For example, Hairol et al. (2015) used Sheridan Gardiner letters and unlimited viewing time, and Ludvigh (1941) used F, E, C, L, T letters and did not specify the viewing time.

**Figure 5.**
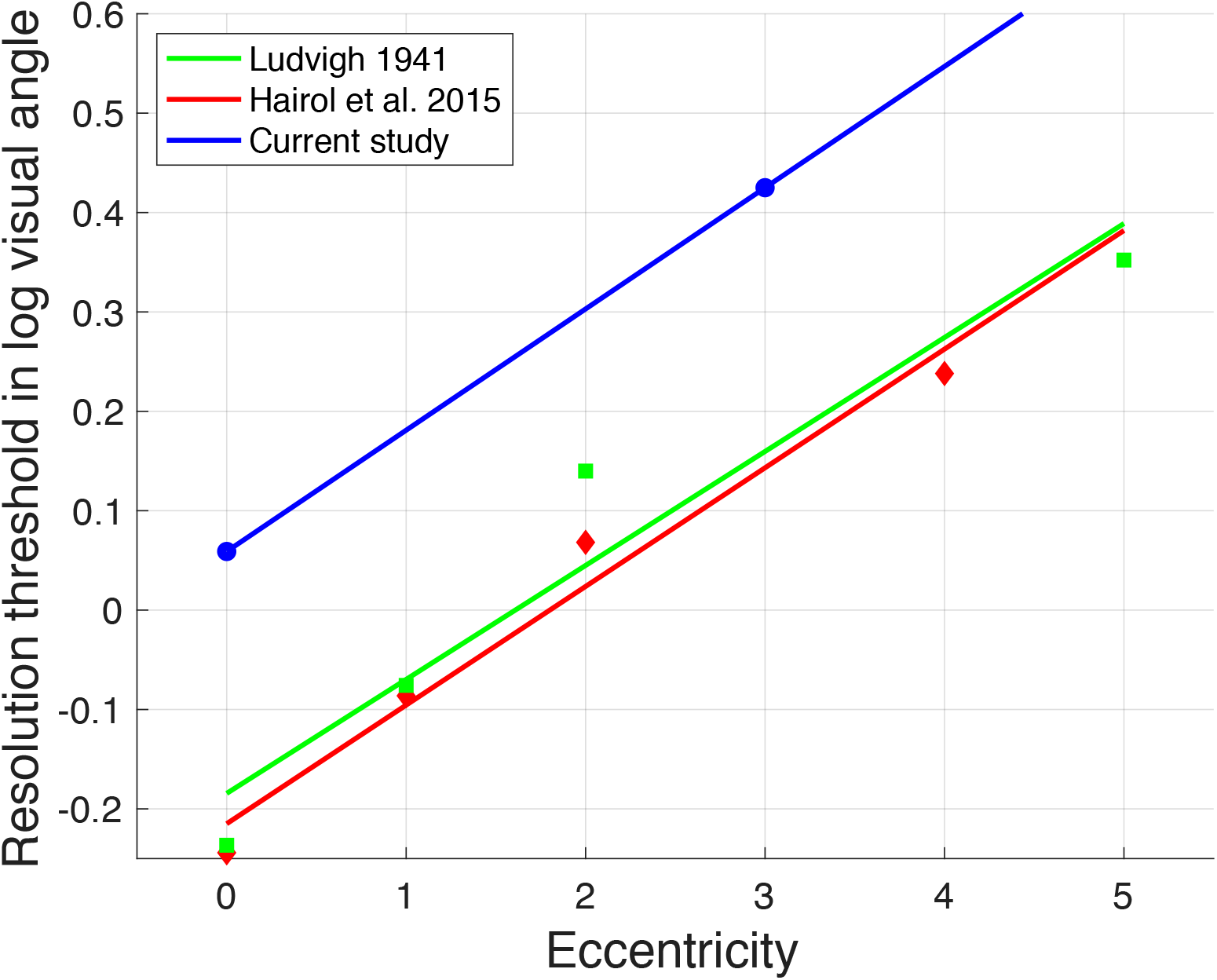
shows the resolution thresholds as a function of eccentricity for two previous studies (green and red) and for the current study (blue).

Similar to previous experiments, our results show significant differences of resolution thresholds of individual Sloan letters (Alexander et al., 1997; Reich & Bedell, 2000; Hamm, Yeoman et al., 2018). Figure 6 shows the resolution thresholds of individual Sloan letters (central) compared to two previous studies (Hamm, 2018; Reich & Bedell, 2000). Our results were similar to Reich & Bedell’s (2000) results and both were different from Hamm’s (2018) results. One might suggest that theses differences in thresholds could be the consequence of employing different psychophysical procedures. Reich & Bedell (2000) and the current experiments employed the method of constant stimuli, whereas Hamm et al. (2018) used a Bayesian adaptive staircase procedure (QUEST). However, the pattern of the differences of letters resolution thresholds were similar in the three studies. This suggests that using different psychophysical procedures to measure the resolution thresholds has no influence on the pattern of the differences of the letter resolution thresholds.

**Figure 6.**
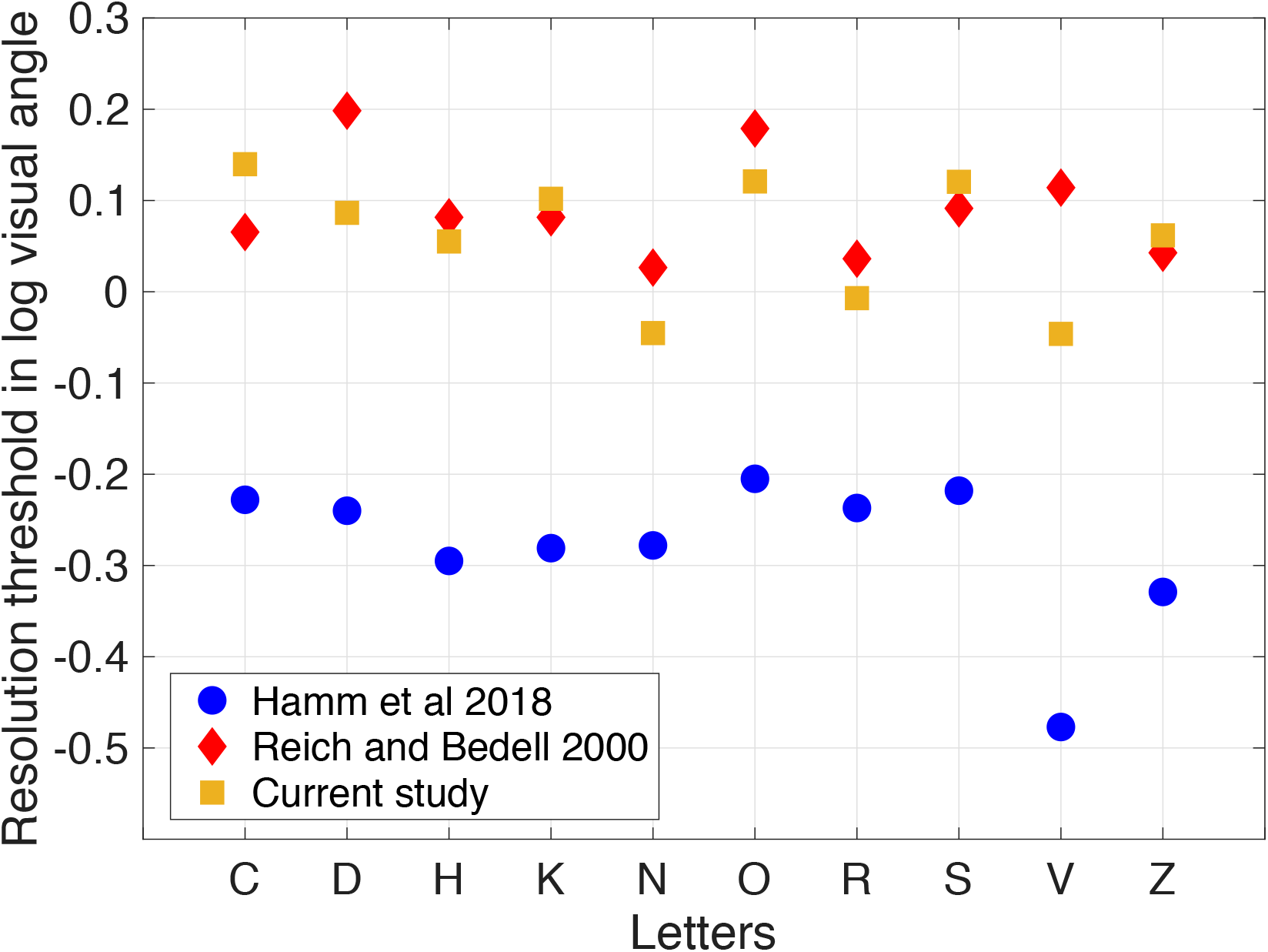
shows the resolution thresholds of individual Sloan letters for the current and two previous studies (Hamm et al., 2018; Reich & Bedell, 2000).

In order to investigate the effect of response biases on the resolution thresholds for individual Sloan letters, Luce’s choice model (Luce, 1963) was used to estimate both the letter similarity and bias parameters for each letter. Theses parameters were used to test their correlation to the differences of the letter resolution thresholds. The top five similarly perceived letters pairs in term of *η* parameter (for example **C** and **O** is similarly perceived letters pair) were compared to what was reported previously (Table 3). Four out of five similarity pairs at the fovea were found to be similar to the results of Hamm et al. (2018), but with different *η* parameter ranks. As mentioned above, this could be the results of employing different methods to determine the thresholds. Four out of five pairs in the fovea and three out of five pairs at the upper paracentral visual field were found to be similar to what was reported by Reich and Bedell (2000). These differences could be the result of using different methods to determine the letters similarity. We used the Luce’s model to determine, whereas Reich and Bedell (2000) used alternative method to determine similarity pairs. Using separate sessions for the fovea and periphery in Reich and Bedell’s (2000) study could be an additional factor; we used interleaved presentations across locations. Furthermore, the superior visual field location in Reich and Bedell’s (2000) study was at 10° compared to 3° in the current experiment. However, independent of this, differences are expected since Reich and Bedell (2000) used 25 alphabet letters. In this case there would be a higher chance to obtain more and different combinations of confusion pairs.

**Table 3.**
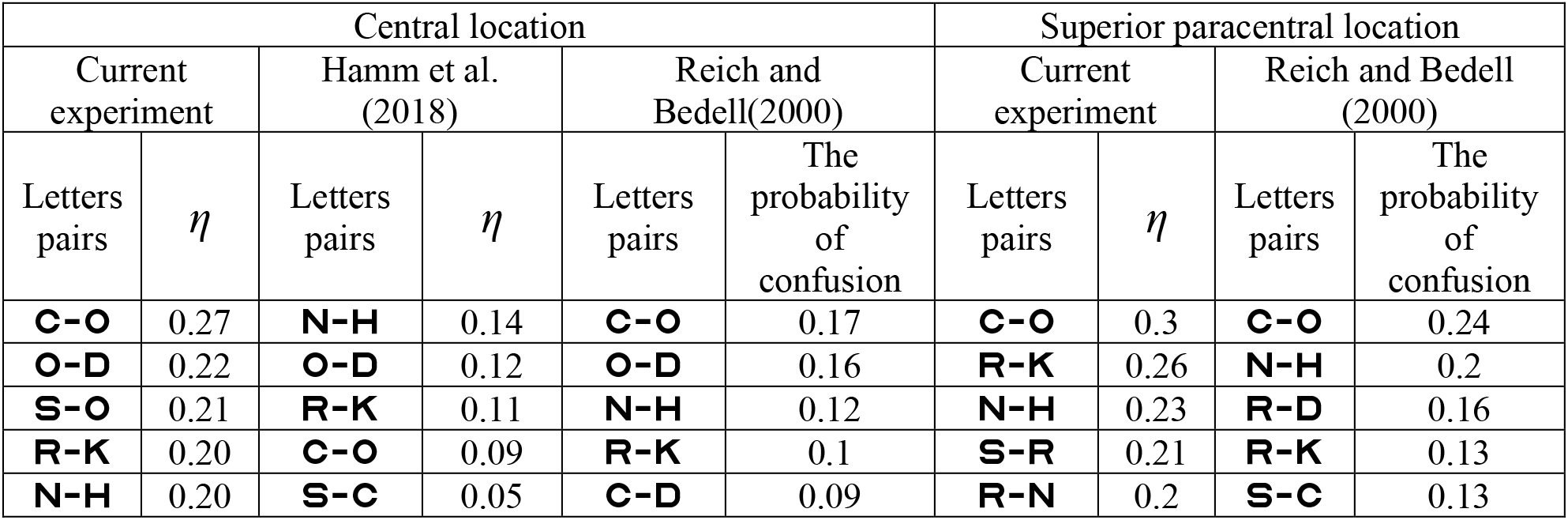
shows the top five similarly perceived letters pairs in the current and two previous studies (Hamm et al., 2018; Reich & Bedell, 2000).

| Central location |  |  |  |  |  | Superior paracentral location |  |  |  |
| --- | --- | --- | --- | --- | --- | --- | --- | --- | --- |
| Current experiment |  | Hamm et al. (2018) |  | Reich and Bedell(2000) |  | Current experiment |  | Reich and Bedell (2000) |  |
| Letters pairs | $\eta$ | Letters pairs | $\eta$ | Letters pairs | The probability of confusion | Letters pairs | $\eta$ | Letters pairs | The probability of confusion |
| **C-O** | 0.27 | **N-H** | 0.14 | **C-O** | 0.17 | **C-O** | 0.3 | **C-O** | 0.24 |
| **O-D** | 0.22 | **O-D** | 0.12 | **O-D** | 0.16 | **R-K** | 0.26 | **N-H** | 0.2 |
| **S-O** | 0.21 | **R-K** | 0.11 | **N-H** | 0.12 | **N-H** | 0.23 | **R-D** | 0.16 |
| **R-K** | 0.20 | **C-O** | 0.09 | **R-K** | 0.1 | **S-R** | 0.21 | **R-K** | 0.13 |
| **N-H** | 0.20 | **S-C** | 0.05 | **C-D** | 0.09 | **R-N** | 0.2 | **S-C** | 0.13 |

The response biases at the central location (**N**, **R**, **V**) were similar (except one letter) to what has been reported by Hamm et al. (2018) (**K**, **R**, **V**). Surprisingly, Pearson product-moment correlation test showed a strong correlation (*r* = 0.74, n = 9,*p* < .05) (excluding the letter **K**) in the response biases between the current study and Hamm et al.’s (2018) experiment. This suggests that the response biases in Sloan letters are consistent when the data are pooled across subjects even when employing different psychophysical methods.

Interestingly, unlike letter similarities, the response biases showed strong correlations to the resolution differences in the current study. In Hamm et al.’s (2018) experiment, the response biases did not explain or had little influence on the thresholds differences. As mentioned above, the reason might be due to the method used to estimate the thresholds. In the method of constant stimuli, the letter sizes at pure guessing levels (sensory-independent, where one would expect more response biases) were larger than the letter sizes at pure guessing level in the QUEST method where the majority of letter sizes would be at the confusion level. This might explain why the response bias did not explain the threshold differences in Hamm et al.’s (2018) study but did in the current study.

In summary, the response biases estimated by Luce choice model (Luce, 1963) revealed a significant effect on the differences of resolution thresholds of individual Sloan letters measured by the method of constant stimuli at the central and paracentral locations. We can therefore conclude that response biases can potentially lead to an overestimation of measured thresholds of the biased letters on the expense of the others.

## Appendix Luce’s choice model

The fitting algorithm was originally presented by Smith (1982). The fitting procedure consists of two steps. The first step is to find the maximum likelihood estimate of the model. The iterative proportional fitting is used to converge the raw (confusion) matrix to the maximum likelihood estimate of the model, as follows: The starting matrix values are all ones. In order to perform the first iteration (cycle), adjustment for rows, columns and similarities are carried out successively. The adjustment for rows is performed by dividing the value of each cell by the sum of the corresponding row values of the starting matrix (ones for the first cycle) then multiply by the marginal sum of the corresponding row values of the raw (confusion) matrix, followed by the adjustment for columns then for similarities. The resulting matrix will be the starting matrix for the second iteration (cycle). These iterations (cycles) are repeated until there is no appreciated change in the estimated values. The resulted matrix is the maximum likelihood estimate of the model. The second step is to compute the parameters of the response bias vector and the similarity matrix from the maximum likelihood estimate of the model, using the Eqs. 3 and 4 respectively.

The MATLAB code to compute the response biases and similarities parameters using Luce’s choice model was written by one of the authors (Hatem Barhoom) and can be downloaded from the link: https://github.com/HBarhoom/Codes-

